# Running Style and Stability During Uphill Running Are Largely Preserved with Increasing Shoe Sole Thickness

**DOI:** 10.64898/2026.04.16.719110

**Authors:** Cagla Kettner, Bernd J. Stetter, Thorsten Stein

## Abstract

Advanced footwear technology (AFT) shoes incorporate increased sole thickness and compliant midsole materials that may alter running biomechanics. While these effects have been widely studied during level running, little is known about how sole thickness influences running style and stability during uphill running. This study examined the effects of two AFT shoes differing in sole thickness (35 mm-AFT35; 50 mm-AFT50) and a traditional control shoe (27 mm-CON27) on running style and stability during uphill running. Seventeen experienced male runners performed treadmill running at a 10% incline at 6.5 and 10 km/h in three shoe conditions. Running style was assessed using duty factor, normalized step frequency, center-of-mass oscillation, vertical and leg stiffness, and lower-limb joint kinematics. Running stability was evaluated using local dynamic stability via the maximum Lyapunov exponent and detrended fluctuation analysis of stride time. Duty factor and normalized step frequency did not differ between shoes. However, AFT shoes showed greater center-of-mass oscillation (p = 0.004), lower vertical stiffness (p = 0.022) compared to CON27. Joint kinematics revealed significant shoe effects at the ankle (p = 0.001), particularly increased dorsiflexion and eversion in AFT conditions. Running stability showed only minor changes. Local dynamic stability differed at the trunk (p = 0.027), with reduced stability in AFT50 compared with CON27 (p = 0.006), while global stability remained unchanged. No shoe × speed interactions were observed for any variable. Overall, uphill running style and stability remained largely preserved across shoe conditions, suggesting that sole thickness alone had limited influence.

## Introduction

Over the past decade, running shoe design has evolved substantially with the introduction of Advanced Footwear Technologies (AFT) [1–3]. These shoes typically combine increased sole thickness with lightweight, compliant, and resilient midsole foams, and integrate carbon plates or rods designed to modify longitudinal bending stiffness and reduce mechanical energy loss at the foot joints [4–6]. In response to their potential performance advantages, World Athletics limited sole thickness (40 mm ≥) and restricted the number of embedded rigid elements [7].

While AFT shoes have been widely studied, the specific biomechanical contribution of sole thickness independent of other shoe features remains unclear. Some studies suggest that thicker soles may enhance performance by effectively increasing leg length and stride length [8,9]. Others argue that performance benefits primarily arise from interactions between compliant foams and stiffening elements rather than sole thickness per se [10,11]. At the same time, concerns have been raised that increased sole thickness could compromise ankle stability and potentially elevate injury risk [9,12]. A recent systematic review concluded that thicker soles tend to increase stance time and ankle dorsiflexion at foot contact, while effects on knee and hip kinematics remain inconsistent [13]. Notably, the influence of sole thickness on overall running style and stability remains insufficiently understood and has primarily been studied in level running and, more recently, during downhill running [14].

Running style is considered an important determinant of both running performance and injury risk [15–17]. A practical framework to operationalize running style was proposed by Van Oeveren et al. (2021), who introduced a dual-axis model based on two main spatiotemporal variables, duty factor (DF) and step frequency normalized to leg length (SF_norm_). This approach allows for an intuitive classification of observable strategies such as “small-step” and “large-step” running and has been applied to quantify style adaptations across different conditions, including shoe interventions [19,20]. However, these spatiotemporal variables alone may not fully reflect the underlying biomechanical mechanisms [14,21,22]. Changes in sole thickness, for example, were shown to particularly influence ankle and knee kinematics during landing and midstance [22]. Moreover, recent findings indicate that sole thickness may affect leg stiffness and vertical center-of-mass oscillation (COM_osc_), suggesting broader implications for running style modulation [23].

Another important component of locomotor control is running stability, which has been associated with both running performance and injury risk [24–26]. In running research, local and global stability are most commonly assessed [27–29]. Local dynamic stability is usually quantified using the maximum Lyapunov exponent (MLE), which reflects sensitivity to small perturbations and has been shown to respond to fatigue, running speed, and skill level [24,25,30]. Global stability, often assessed using detrended fluctuation analysis (DFA) of stride time, provides insight into long-range temporal correlations. Thereby, stronger correlations are interpreted as reduced adaptability and thus lower global stability [28,29,31]. Despite the relevance of these measures, relatively few studies have examined how shoe properties such as sole thickness influence running stability.

Importantly, nearly all previous studies examining sole thickness have focused on level running [13]. Recent work has begun to explore the influence of sole thickness under other locomotor conditions, such as downhill running [14]. Parts of the conceptual framework presented in the current study build upon this previous investigation. However, uphill running represents a fundamentally different locomotor task. Compared with level running, uphill locomotion requires greater positive mechanical work to elevate the body’s center-of-mass, leading to higher propulsive demands and altered joint work distribution [32,33]. Runners typically respond to these demands by increasing cadence, shortening step length, and increasing DF [34,35]. At the same time, impact-related braking forces are reduced and muscular work becomes more concentric in nature [35,36]. Because of these task-specific mechanical requirements, shoe features that are shown to be beneficial during steady-state level running, such as highly resilient foams optimized for elastic energy return [37], may exert different effects during uphill running [38].

Despite the practical importance of uphill running in disciplines such as trail running and mountain racing, the isolated effects of shoe sole thickness under uphill conditions have not yet been systematically examined. Previous studies investigating shoe effects during uphill running have typically compared shoe models differing in multiple design characteristics, making it difficult to attribute observed effects specifically to sole thickness [38–40]. A better understanding of how sole thickness influences running mechanics during uphill locomotion may therefore provide valuable insights for both shoe design and injury prevention.

Accordingly, the present study investigated the effects of three different shoes (two AFT shoes differing mainly in sole thickness [35 mm and 50 mm], and one traditional shoe [27 mm]) on running style and stability during uphill running at two speeds. It was hypothesized that AFT shoes with thicker soles (35 and 50 mm) would modulate running style and reduce both local and global stability compared with a traditional thinner shoe (27 mm). Second, the shoe with the greatest stack height (50 mm) was expected to induce greater changes in running style and further reductions in stability compared with the moderately thick AFT shoe (35 mm). Third, these shoe-related effects were hypothesized to be pronounced at the higher running speed.

## Methods

### Participants

Seventeen healthy, experienced male runners participated in the study (age: 25.7 ± 3.9 years; height: 1.77 ± 0.04 m; body mass: 68.1 ± 6.0 kg). All participants were regularly active runners (4.2 ± 1.8 sessions per week; 33.7 ± 22.4 km per week). Only male participants were included because standardized test shoes were available in a single size (US men’s 9). Sample size was determined based on previous studies with comparable designs [41–43], and expected medium effect sizes (effect size, f = 0.25; power 0.80). The recruitment period for this study was between 17.07.2023–05.12.2023. All participants provided written informed consent. The study was approved by the Ethics Committee of the Karlsruhe Institute of Technology.

### Shoes

Three running shoes were tested: two AFT models (AFT50 and AFT35) and one traditional control shoe (CON27). Heel and forefoot heights were 50 mm and 43 mm for AFT50, 35 mm and 28 mm for AFT35, and 27 mm and 19 mm for CON27. Heel-to-toe drop was similar across conditions (7 mm for AFT50 and AFT35; 8 mm for CON27), and shoe mass was comparable (268 g, 220 g, and 219 g for AFT50, AFT35, and CON27, respectively; all values refer to US men’s size 9). The two AFT models incorporated carbon-based propulsion elements, whereas the control shoe represented a conventional design [14]. To minimize confounding effects, shoes were matched as closely as possible in mass (< 50 g difference) and heel-to-toe drop (7–8 mm). Apart from sole thickness, the AFT models were identical in construction, enabling isolation of thickness-related effects.

### Study design

All measurements were performed on a motorized treadmill (h/p/cosmos Saturn, Nussdorf-Traunstein, Germany). Participants were secured with an overhead safety harness throughout all trials to prevent injury without providing bodyweight support. Participants first completed a 5-min familiarization in their own shoes at a self-selected speed under three slope conditions (0%, 10% downhill, 10% uphill). They then performed three counterbalanced measurement blocks, each corresponding to one standardized shoe condition. Each block began with a 5-min shoe-specific familiarization [44], consisting of 3 min running and 2 min walking at self-selected speed under level condition.

Following familiarization, participants completed trials under three slope conditions: 0% (level), −10% (downhill), and +10% (uphill). The present analysis focuses exclusively on uphill running as each slope represents a distinct motor task with specific biomechanical and energetic characteristics [38,45] and speeds were not metabolically matched across gradients [39]. Results for level [23,46] and downhill [14] conditions have been reported in previous studies. Uphill trials were performed at 6.5 and 10 km/h on a 10% incline, based on prior literature and pilot testing [45]. Participants ran 20–30 s before data acquisition to ensure steady-state speed, followed by 90 s of recording. Recovery periods included 1-min walking between speeds, 2-min standing between slope conditions, and 5-min seated rest between shoes. Perceived exertion was assessed using the Borg scale (6–20; Borg, 1982), and trials proceeded only when ratings were ≤ 12 to ensure that participants performed the trials under non-fatigued conditions.

### Data acquisition and preprocessing

Three-dimensional full-body kinematics were captured using a 16-camera motion capture system (Vicon Motion Systems, Oxford, UK) operating at 200 Hz and 49 reflective markers. Marker trajectories were reconstructed using Vicon Nexus software (v2.15) and subsequently exported for further processing in MATLAB (R2024a; MathWorks Inc., Natick, MA, USA). All trajectories were filtered using a fourth-order low-pass Butterworth filter with a cutoff frequency of 10 Hz [48].

### Data analysis

Consistent with the data acquisition and preprocessing procedures described above, data analysis followed previously established methods [14], with the main steps outlined below. Running style was characterized using spatiotemporal parameters, center-of-mass motion, stiffness variables, and lower-limb joint kinematics. Spatiotemporal measures included DF and SF_norm_. DF was defined as the ratio of stance time to stride time, whereas SF_norm_ was calculated by normalizing step frequency to leg length (greater trochanter to ground distance) and gravitational acceleration following established scaling principles [18,49,50].

Gait events were identified from kinematic data. Initial contact was defined as the instant corresponding to the minimum of the combined heel and toe marker velocities [51], while toe-off was determined as the time point of maximal knee extension in the sagittal plane [52]. These approaches have previously demonstrated acceptable temporal accuracy during treadmill running, with errors typically below 20 ms [51–53], and were therefore considered appropriate for the present analysis.

Joint kinematics and center-of-mass trajectories were obtained using inverse kinematics in OpenSim based on a full-body musculoskeletal model [54]. Each model was individually scaled to minimize marker reconstruction error. COM_osc_ during stance was extracted, and vertical stiffness (k_ver_) as well as leg stiffness (k_leg_) were estimated using spring-mass model formulations that incorporate body mass, running velocity, and temporal gait characteristics [49]. Lower-limb joint angles (ankle, knee, and hip) in the sagittal and frontal planes were time-normalized to the stance phase (101 points) to enable comparisons across conditions.

Running stability was assessed using nonlinear time-series methods. Local dynamic stability was quantified via the MLE computed from segmental kinematic trajectories in vertical direction [27,55]. For each condition, time series comprising 100 consecutive strides were resampled to a fixed length [56] and reconstructed in state space using time-delay embedding. The embedding dimension was determined using the false nearest neighbor method, and the time delay was selected based on the first minimum of the average mutual information function [57]. Divergence of neighboring trajectories was calculated using Rosenstein’s algorithm [58], and the short-term slope of the divergence curve was used as the MLE, with lower values indicating greater local stability.

Global stability was evaluated using DFA of stride time intervals. The stride time series was demeaned and cumulatively integrated, then segmented into windows of varying lengths. Within each window, local trends were removed using polynomial detrending, and root-mean-square fluctuations were calculated. The scaling exponent (DFA-α) was obtained from the slope of the log–log relationship between fluctuation magnitude and window size, where higher values indicate more persistent correlations and reduced global stability [28,31].

### Statistics

Statistical analyses were performed using SPSS (Version 29.0; IBM Corp., Armonk, NY, USA). For variables derived from linear analyses (DF, SF_norm_, COM_osc_, k_ver_, k_leg_), mean values across 25 strides per condition were calculated. Nonlinear measures (MLE and DFA-α) were obtained as single values from time series comprising 100 consecutive strides. Data distribution and sphericity assumptions were evaluated using Kolmogorov–Smirnov and Mauchly’s tests, respectively. When violations of sphericity were detected, Greenhouse–Geisser corrections were applied.

To examine the effects of shoe condition (AFT50, AFT35, CON27) and running speed (6.5 and 10 km/h), two-way repeated-measures ANOVAs were conducted. Effect sizes were reported as partial eta squared (𝜂^2^), with thresholds of ≤ 0.06 (small), 0.06–0.14 (medium), and ≥ 0.14 (large). In the presence of significant main effects or interactions involving shoe condition, post hoc comparisons were performed using paired t-tests with Bonferroni–Holm adjustment. Effect sizes for pairwise comparisons were expressed as Cohen’s d (small ≤ 0.50, medium 0.50–0.80, large ≥ 0.80; Cohen, 1988).

Joint angle time series were analyzed using statistical parametric mapping (SPM; spm1d toolbox) implemented in MATLAB. Normality of the trajectories was assessed within the SPM framework; if violated, nonparametric inference based on permutation testing (1000 iterations) was applied. When significant effects were detected, post hoc paired t-tests were conducted. Reported SPM cluster intervals are provided descriptively, as statistical inference applies to the full time-normalized trajectory rather than discrete time points (Honert & Pataky, 2021).

Analyses of λ_foot_, λ_lower-leg_, λ_upper-leg,_ and joint angle time series were performed separately for both legs. As results were comparable and showed identical significance patterns, only left-leg data are presented. For all statistical analysis, the significance level was set at α = 0.05. Primary focus was placed on main effects of shoe and shoe × speed interactions. Although speed effects are reported for completeness, they were not interpreted, as they were not central to the study hypotheses.

## Results

### Running style

Discrete running style results are shown in Table 1. SF_norm_ and DF did not differ between shoes (Fig 1). COM_osc_ (p = 0.004, 𝜂^2^ = 0.288) and k_ver_ (p = 0.022, 𝜂^2^ = 0.212) differed between shoes. Post-hoc tests showed that COM_osc_ was lower with CON27 compared to AFT50 (p = 0.006, |d| = 0.785) and AFT35 (p = 0.006, |d| = 0.809). In parallel, k_ver_ was higher with CON27 compared to AFT50 (p = 0.027, |d| = 0.640) and AFT35 (p = 0.028, |d| = 0.588). No shoe effects were observed for k_leg_. No interaction effects between shoe and speed were observed for any discrete running style variable.

**Fig 1.**
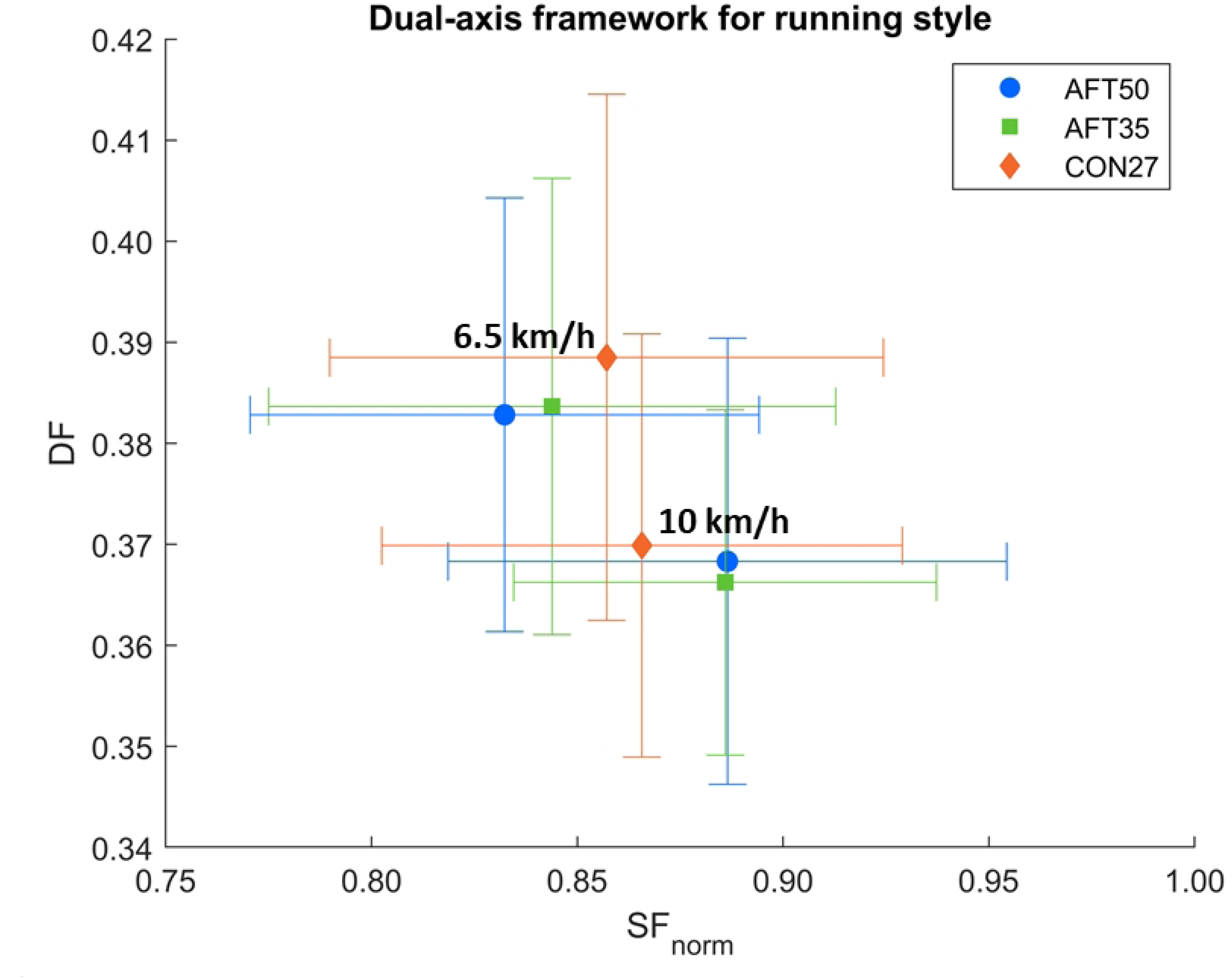
Running style analysis based on the dual-axis framework (van Oeveren et al., 2021) across different shoe conditions [AFT50 (50 mm), AFT35 (35 mm), and CON27 (27 mm)] and running speeds (6.5 and 10 km/h). Symbols represent mean values for each shoe, with error bars indicating standard deviations.

**Table 1.** Variables included in the running style analysis comprised step frequency normalized to leg length (SF_norm_), duty factor (DF), vertical center-of-mass oscillation (COM_osc_), vertical stiffness (k_ver_), and leg stiffness (k_leg_).

| $SF_{norm}$ | | | | | | |
| --- | --- | --- | --- | --- | --- | --- |
| | 6.5 km/h | 10 km/h | rmANOVA | F | p | $\eta_p^2$ |
| AFT50 | $0.84 \pm 0.07$ | $0.89 \pm 0.07$ | Shoe | 2.392 | 0.129 | 0.130 |
| AFT35 | $0.83 \pm 0.06$ | $0.86 \pm 0.06$ | Speed | **27.213** | **< 0.001** | **0.630** |
| CON27 | $0.86 \pm 0.07$ | $0.89 \pm 0.05$ | Interaction | 0.859 | 0.433 | 0.051 |
| DF |  |  |  |  |  |  |
| | 6.5 km/h | 10 km/h | rmANOVA | F | p | $\eta_p^2$ |
| AFT50 | $0.38 \pm 0.02$ | $0.37 \pm 0.02$ | Shoe | 0.266 | 0.768 | 0.016 |
| AFT35 | $0.38 \pm 0.02$ | $0.37 \pm 0.02$ | Speed | **28.430** | **< 0.001** | **0.640** |
| CON27 | $0.39 \pm 0.03$ | $0.37 \pm 0.02$ | Interaction | 1.408 | 0.259 | 0.081 |
| $COM_{osc}$ (cm) | | | | | | |
| | 6.5 km/h | 10 km/h | rmANOVA | F | p | $\eta_p^2$ |
| AFT50 | $7.51 \pm 0.98$ | $7.90 \pm 1.06$ | Shoe | **6.461** | **0.004** | **0.288** |
| AFT35 | $7.67 \pm 1.12$ | $7.95 \pm 1.12$ | Speed | **5.667** | **0.030** | **0.262** |
| CON27 | $7.13 \pm 0.82$ | $7.67 \pm 1.07$ | Interaction | 2.090 | 0.140 | 0.115 |
|  |  |  | *Post-hoc* | t | p | \|d\| |
|  |  |  | AFT50-AFT35 | -0.826 | 0.210 | 0.200 |
|  |  |  | AFT50-CON27 | **3.236** | **0.006** | **0.785** |
|  |  |  | AFT35-CON27 | **3.337** | **0.006** | **0.809** |
| k<sub>ver</sub> (kN/m) |  |  |  |  |  |  |
| | 6.5 km/h | 10 km/h | rmANOVA | F | p | $\eta_p^2$ |
| AFT50 | 16.00 ±<br>2.40 | 16.23 ± 2.31 | Shoe | **4.312** | **0.022** | **0.212** |
| AFT35 | 15.99 ±<br>2.92 | 15.98 ± 2.58 | Speed | 0.171 | 0.684 | 0.011 |
| CON27 | 16.69 ±<br>2.80 | 16.52 ± 2.40 | Interaction | 0.166 | 0.848 | 0.010 |
|  |  |  | *Post-hoc* | t | p | \|d\| |
|  |  |  | AFT50-AFT35 | 0.594 | 0.280 | 0.144 |
|  |  |  | AFT50-CON27 | **-2.638** | **0.027** | **0.640** |
|  |  |  | AFT35-CON27 | **-2.423** | **0.028** | **0.588** |
| k<sub>leg</sub> (kN/m) |  |  |  |  |  |  |
| | 6.5 km/h | 10 km/h | rmANOVA | F | p | $\eta_p^2$ |
| AFT50 | 8.05 ± 1.52 | 5.46 ± 1.11 | Shoe | 1.080 | 0.331 | 0.063 |
| AFT35 | 7.87 ± 1.70 | 5.36 ± 1.16 | Speed | **119.086** | **< 0.001** | **0.882** |
| CON27 | 8.10 ± 1.87 | 5.66 ± 1.02 | Interaction | 0.157 | 0.788 | 0.010 |

Shoe sole thickness conditions are denoted as AFT50 (50 mm), AFT35 (35 mm), and CON27 (27 mm). Data are presented as mean ± standard deviation. Post hoc comparisons were performed on mean values at 6.5 and 10 km/h. Statistically significant effects are highlighted in **bold**.

Joint angle time-series results are illustrated in Figs 2 and 3 for the sagittal and frontal planes, respectively. SPM analysis revealed significant shoe effects for the sagittal ankle angle (p = 0.001), with a cluster spanning the entire stance phase (0–100%; Fig 2). Post-hoc comparisons indicated higher dorsiflexion in AFT35 compared with CON27 (p = 0.002; 7–55% of stance; Fig S1).

**Fig 2.**
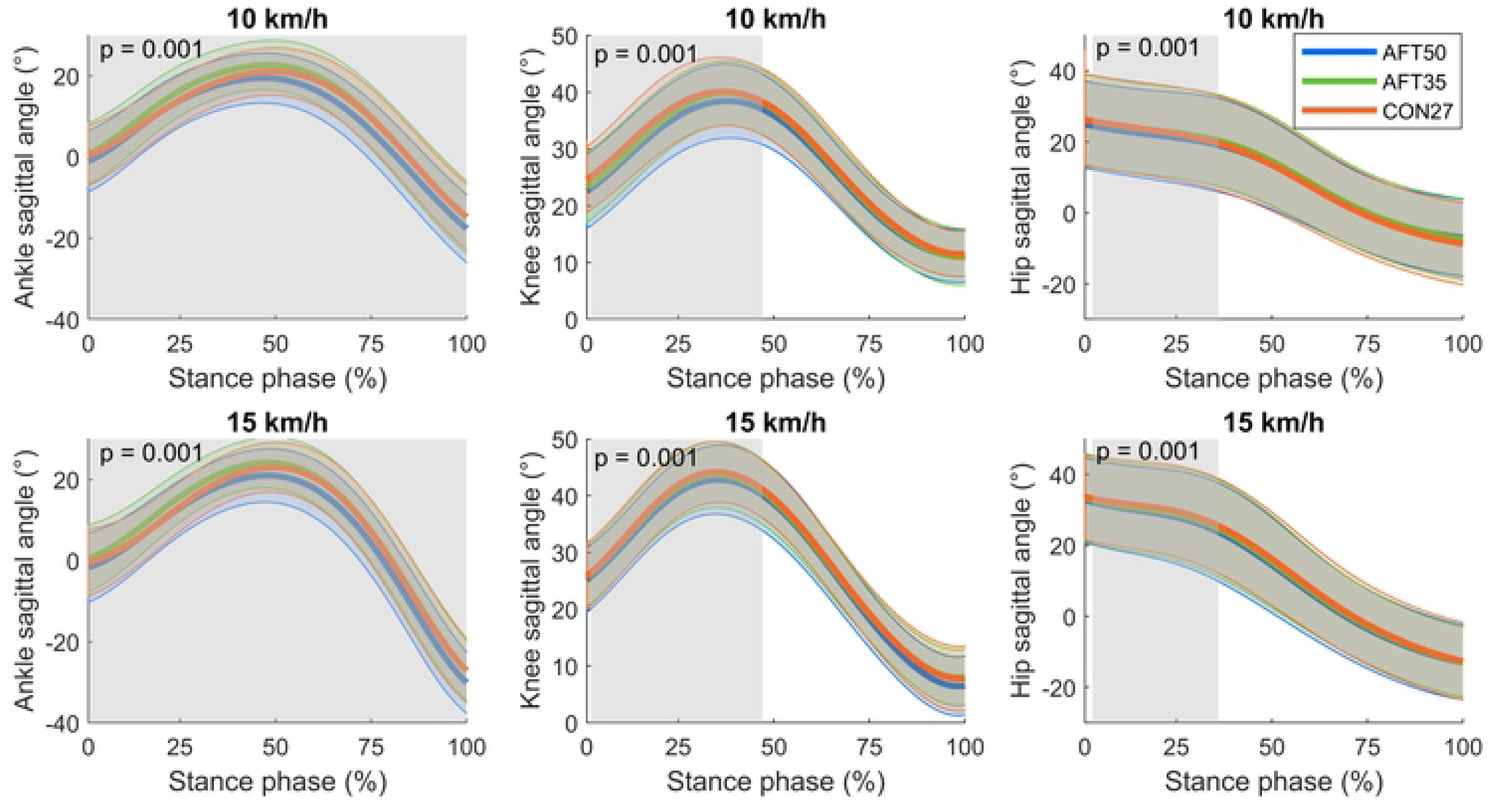
Sagittal-plane joint angle trajectories of the ankle, knee, and hip presented as mean curves (thick lines) with ± standard deviation (thin lines). Positive values denote joint flexion (ankle: dorsiflexion). Grey shaded rectangles indicate significant shoe effects independent of running speed, with corresponding cluster-level p-values reported. Shoe conditions are abbreviated as AFT50 (50 mm), AFT35 (35 mm), and CON27 (27 mm).

**Fig 3.**
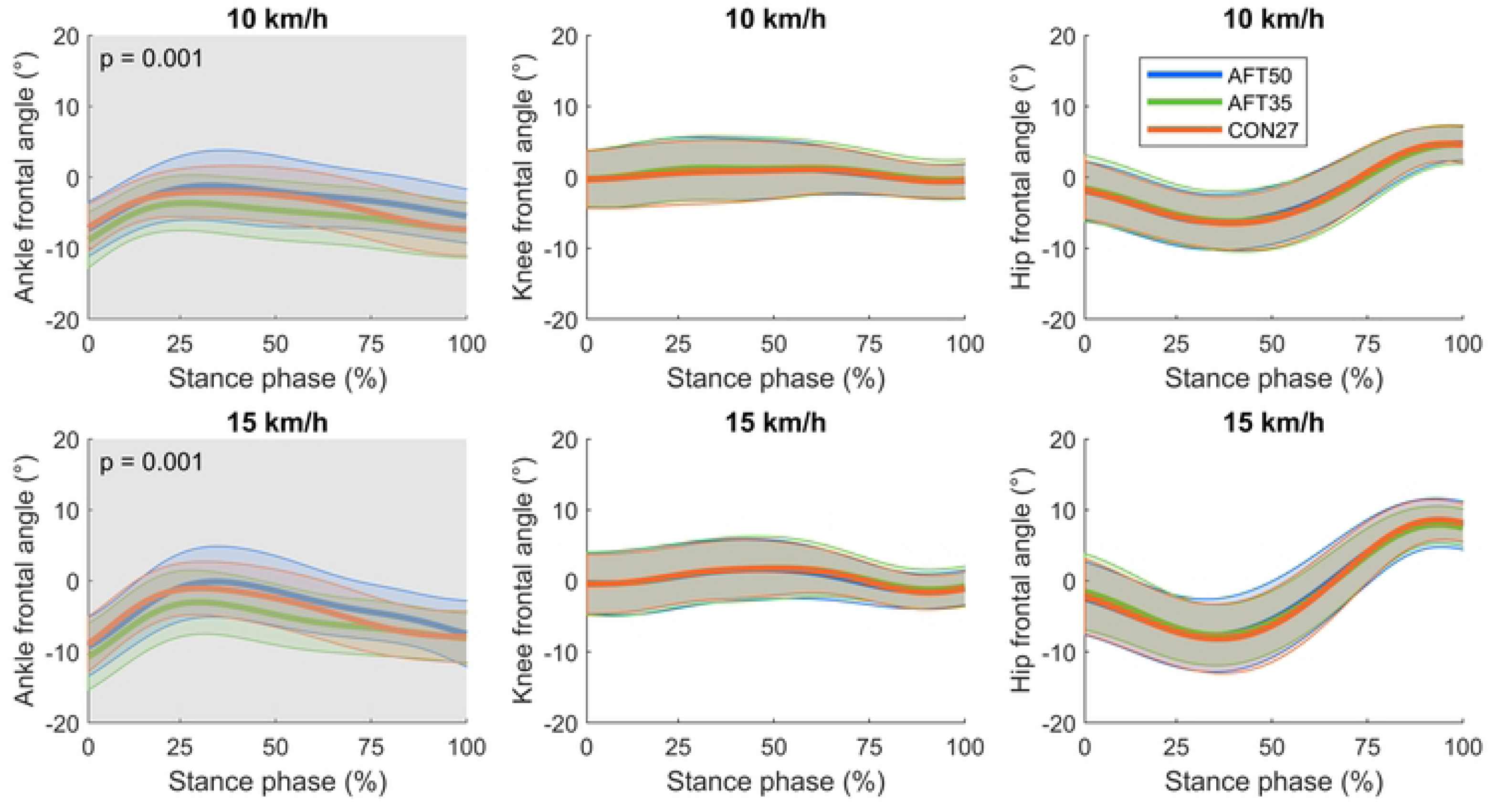
Frontal-plane joint angle trajectories of the ankle, knee, and hip presented as mean curves (thick lines) with ± standard deviation (thin lines). Positive values denote joint abduction at the knee and hip and eversion at the ankle. Grey shaded rectangles indicate significant shoe effects independent of running speed, with corresponding cluster-level p-values reported. Shoe conditions are abbreviated as AFT50 (50 mm), AFT35 (35 mm), and CON27 (27 mm).

A significant shoe effect was observed for the sagittal knee angle (p = 0.001; 2–45% of stance; Fig 2). However, post-hoc analysis revealed no significant pairwise shoe differences (Fig S2). The sagittal hip angle likewise showed a significant effect of shoe (p = 0.001; 1–36% of stance; Fig 2), although post-hoc analysis again revealed no significant pairwise shoe differences (Fig S3).

In the frontal plane, significant differences between shoes were detected only for the ankle angle (p = 0.001), with a cluster spanning the entire stance phase (0–100%; Fig 3). Specifically, AFT50 exhibited greater eversion than AFT35 (p < 0.001; 0–100% of stance) and CON27 (p = 0.001; 63–100% of stance; Fig S3). No interaction effects between shoe and speed were observed for any joint angle time series.

### Running stability

Running stability results are presented in Table 2. Local dynamic stability differed between shoes only at the trunk (p = 0.027, 𝜂^2^ = 0.202). Post-hoc analysis revealed that AFT50 resulted in a higher λ_trunk_, indicating lower local trunk stability, compared with CON27 (p = 0.006, |d| = 0.811). No other local stability measures differed between shoes. Global stability did not differ between shoe conditions. No shoe–speed interaction effects were observed for any stability variable.

**Table 2.** Variables included in the running stability analysis comprised local dynamic stability of the foot (λ_foot_), lower-leg (λ_lower-leg_), upper-leg (λ_upper-leg_), hip (λ_hip_), and trunk (λ_trunk_), detrended fluctuation scaling component of stride time (DFA-α), peak eversion (MAX_eversion_), and eversion duration (t_eversion_) during stance.

| $\lambda_{\text{foot}}$ | | | | | | |
| --- | --- | --- | --- | --- | --- | --- |
| | 6.5 km/h | 10 km/h | rmANOVA | F | p | $\eta_p^2$ |
| AFT50 | $0.86 \pm 0.11$ | $0.94 \pm 0.09$ | Shoe | 0.766 | 0.473 | 0.046 |
| AFT35 | $0.88 \pm 0.13$ | $0.94 \pm 0.10$ | Speed | **19.267** | **<0.001** | **0.546** |
| CON27 | $0.85 \pm 0.10$ | $0.92 \pm 0.10$ | Interaction | 0.222 | 0.395 | 0.014 |
| $\lambda_{\text{lower-leg}}$ | | | | | | |
| | 6.5 km/h | 10 km/h | rmANOVA | F | p | $\eta_p^2$ |
| AFT50 | $0.44 \pm 0.11$ | $0.55 \pm 0.12$ | Shoe | 1.660 | 0.206 | 0.094 |
| AFT35 | $0.48 \pm 0.08$ | $0.55 \pm 0.06$ | Speed | **42.520** | **<0.001** | **0.727** |
| CON27 | $0.45 \pm 0.09$ | $0.51 \pm 0.12$ | Interaction | 1.247 | 0.301 | 0.072 |

| $\lambda_{\text{upper-leg}}$ | | | | | | |
| --- | --- | --- | --- | --- | --- | --- |
| | 6.5 km/h | 10 km/h | rmANOVA | F | p | $\eta_p^2$ |
| AFT50 | $1.29 \pm 0.13$ | $1.26 \pm 0.12$ | Shoe | 2.615 | 0.089 | 0.140 |
| AFT35 | $1.25 \pm 0.12$ | $1.23 \pm 0.12$ | Speed | 2.302 | 0.149 | 0.126 |
| CON27 | $1.23 \pm 0.14$ | $1.21 \pm 0.11$ | Interaction | 0.100 | 0.905 | 0.006 |
| $\lambda_{\text{hip}}$ | | | | | | |
| | 6.5 km/h | 10 km/h | rmANOVA | F | p | $\eta_p^2$ |
| AFT50 | $1.35 \pm 0.12$ | $1.41 \pm 0.16$ | Shoe | 1.734 | 0.193 | 0.098 |
| AFT35 | $1.34 \pm 0.18$ | $1.42 \pm 0.17$ | Speed | **6.318** | **0.023** | **0.283** |
| CON27 | $1.31 \pm 0.14$ | $1.36 \pm 0.14$ | Interaction | 0.174 | 0.841 | 0.011 |
| $\lambda_{\text{trunk}}$ | | | | | | |
| | 6.5 km/h | 10 km/h | rmANOVA | F | p | $\eta_p^2$ |
| AFT50 | $1.02 \pm 0.14$ | $1.09 \pm 0.10$ | Shoe | **4.051** | **0.027** | **0.202** |
| AFT35 | $1.02 \pm 0.11$ | $1.07 \pm 0.12$ | Speed | **6.593** | **0.021** | **0.292** |
| CON27 | $1.00 \pm 0.12$ | $1.00 \pm 0.16$ | Interaction | 1.058 | 0.359 | 0.062 |
|  |  |  | *Post-hoc* | t | p | \|d\| |
|  |  |  | AFT50-AFT35 | 0.558 | 0.292 | 1.135 |
|  |  |  | AFT50-CON27 | **3.343** | **0.006** | **0.811** |
|  |  |  | AFT35-CON27 | 1.930 | 0.072 | 0.087 |
| DFA-α | | | | | | |
| | 6.5 km/h | 10 km/h | rmANOVA | F | p | $\eta_p^2$ |
| AFT50 | $0.56 \pm 0.17$ | $0.61 \pm 0.15$ | Shoe | 0.063 | 0.893 | 0.004 |
| AFT35 | 0.55 ± 0.21 | 0.64 ± 0.15 | Speed | **4.602** | **0.048** | **0.223** |
| CON27 | 0.55 ± 0.14 | 0.63 ± 0.16 | Interaction | 0.218 | 0.731 | 0.02 |

Shoe sole thickness conditions are denoted as AFT50 (50 mm), AFT35 (35 mm), and CON27 (27 mm). Data are presented as mean ± standard deviation. Post hoc comparisons were performed on mean values at 6.5 and 10 km/h. Statistically significant effects are highlighted in **bold**.

## Discussion

The purpose of this study was to investigate the effects of different shoe sole thicknesses on running style and running stability during uphill running. Overall, AFT shoes influenced vertical center-of-mass motion, vertical stiffness, and ankle joint kinematics compared with the traditional shoe, while the spatiotemporal measures of running style remained unchanged. Differences between the two AFT models were limited to frontal ankle kinematics, suggesting that sole thickness alone had a limited influence on running style under uphill conditions. Running stability showed only a small reduction in trunk local stability in the thickest AFT shoe compared with the traditional shoe. Moreover, the effects of shoes were consistent across the two running speeds, as no interactions between shoe and running speed were observed for either running style or stability measures.

### Spatiotemporal running style remained unchanged despite altered center-of-mass motion and ankle kinematics

The first hypothesis proposed that thicker AFT shoes would modulate running style compared with the traditional shoe. The results provided partial support for this assumption. While COM_osc_ and k_ver_ differed between AFT shoes and the traditional shoe, the key spatiotemporal descriptors of running style within the dual-axis framework (SF_norm_ and DF) remained unchanged.

The absence of shoe effects for SF_norm_ and DF suggests that runners maintained a consistent spatiotemporal pattern across shoe conditions. This may partly relate to the task-specific mechanical demands of uphill running. Compared with level running, uphill locomotion requires greater net positive mechanical work to elevate the center-of-mass, thereby increasing propulsive demands and altering the distribution of joint work across the lower limb [32,33]. Runners typically respond by adopting shorter step lengths together with higher cadence and duty factors to maintain forward progression against gravity [34,35]. At the same time, impact-related loading and eccentric braking demands are reduced, increasing reliance on concentric muscle work [35,36]. Under these conditions, locomotion may depend less on elastic energy storage and return than during level running. Consequently, shoe characteristics that influence elastic rebound, such as compliant and resilient midsoles, may exert smaller effects on the spatiotemporal characteristics of the stride, while still affecting selected mechanical or kinematic variables.

The slightly greater COM_osc_ (< 1 cm) observed in the AFT shoes indicates a small reduction in vertical system stiffness compared with the traditional shoe. Because k_ver_ is inversely related to center-of-mass displacement within the spring–mass framework, the lower k_ver_ values observed in the AFT shoes are consistent with the higher COM_osc_ [49,61]. Joint kinematics did not show consistent increases in lower-limb flexion that would clearly explain this difference, suggesting that the changes in center-of-mass motion may result from subtle adjustments in whole-body mechanics or interactions between the runner and the shoe–surface interface rather than from large joint-level adaptations.

Joint kinematics offered further insight into how runners adapted to the different shoe conditions. In the sagittal plane, AFT35 exhibited greater ankle dorsiflexion compared with CON27. Previous studies have likewise reported increased ankle dorsiflexion with greater shoe sole thickness, suggesting that sole thickness may influence ankle joint mechanics during the stance phase of running [13,14,22]. Although the SPM analysis also indicated overall shoe effects at the knee and hip, no pairwise differences between shoes were identified, suggesting that these effects were small. This indicates that the primary kinematic adaptations occurred at the foot–ankle complex, which directly interacts with the shoe. In the frontal plane, the thickest shoe exhibited greater ankle eversion compared with both AFT35 and CON27. Increased ankle eversion with thicker shoe soles has been reported previously and is often discussed in the context of potential stability challenges associated with thicker soles [12,22]. A higher platform may increase frontal-plane ankle motion even when overall joint kinematics remains largely unchanged. Together, these findings suggest that shoe-related biomechanical adaptations during uphill running were primarily observed at the ankle joint.

Importantly, these differences primarily emerged when comparing the AFT shoes with the traditional shoe, whereas differences between the two AFT models were limited. Because AFT35 and AFT50 share similar design characteristics aside from sole thickness, the comparisons between these shoes more directly reflect the isolated effect of sole thickness. In contrast, comparisons between the AFT shoes and CON27 likely reflect the combined influence of several shoe properties associated with AFT designs, such as midsole compliance, energy return, or shoe geometry [5,6]. The largely similar responses between AFT35 and AFT50 suggest that sole thickness alone had only a limited influence on running style during uphill running, which does not support the second hypothesis. An exception was observed in the frontal plane, where the thickest shoe exhibited greater ankle eversion than both AFT35 and CON27. Increased eversion with thicker soles has previously been attributed to the larger lever arm between the foot and the ground when the foot is positioned further above the contact surface [12,22]. A higher platform may therefore modulate frontal-plane ankle motion even when the overall running style remain largely unchanged. The absence of consistent differences at the knee and hip suggests that shoe-related effects remained primarily localized at the foot–ankle complex, particularly in the frontal plane.

Finally, no interactions between shoe and running speed were observed, indicating that the biomechanical differences between shoes were consistent across the two running speeds. This finding does not support the hypothesis that shoe-related effects would become more pronounced at higher speeds. One possible explanation is that the biomechanical demands of uphill running are primarily governed by the requirement to generate positive mechanical work to elevate the center-of-mass [32,33]. As a result, runners may adopt similar movement strategies across moderate speed changes in order to maintain forward progression against gravity. Under these conditions, shoe-related adaptations may remain consistent across speeds.

Overall, the findings indicate that AFT shoes influenced vertical center-of-mass motion, vertical stiffness, and ankle joint kinematics during uphill running, while the spatiotemporal running style defined by DF and SF_norm_ remained unchanged. Moreover, the limited differences between the two AFT shoes suggest that sole thickness alone played a relatively minor role, whereas other design characteristics of AFT shoes may have contributed more strongly to the observed biomechanical adaptations.

### Running stability remained largely unchanged across shoe conditions

Partially in line with the first hypothesis, trunk local stability was reduced with the thickest AFT shoe compared with the traditional shoe as reflected by the higher λ_trunk_. This finding suggests that the shoe with the greatest sole thickness reduced the ability of the trunk segment to compensate for small perturbations during uphill running. In contrast, no other segmental local stability measures differed between shoes. Moreover, global stability, assessed with DFA of stride time, remained unaffected.

The largely unchanged stability outcomes indicate that increased sole thickness alone does not substantially compromise running stability during uphill locomotion. This finding contradicts the initial expectation that thicker AFT soles would reduce both local and global stability.

Instead, the present results suggest that the locomotor system can maintain stability despite variations in shoes under uphill conditions. Similar observations have been reported in previous running studies where local dynamic stability remained largely unchanged across shoe interventions, indicating that runners may rapidly adapt neuromuscular control strategies to maintain stable movement patterns [14,23,24]

The reduction in trunk stability observed with the thickest shoe compared with the traditional shoe indicates that increased sole thickness may influence trunk stability during uphill running. However, because trunk stability did not differ between the two AFT shoes, these findings do not support the second hypothesis that greater sole thickness would induce larger reductions in stability. This pattern suggests that the observed biomechanical effects cannot be explained by sole thickness alone. Instead, interactions with other shoe properties, such as midsole compliance or longitudinal bending stiffness, may contribute to the biomechanical responses observed when running with AFT shoes [5,6]. The absence of corresponding changes in lower-limb segment stability further suggests that any destabilizing influence was small and segment-specific rather than reflecting a global alteration of running control.

Global stability, reflected by stride-to-stride temporal organization, did not differ between shoe conditions. DFA-α values therefore indicate that the long-range correlations of stride timing were preserved across shoe conditions, suggesting unchanged adaptability of the locomotor system [28,31].

Finally, no interaction between shoe and running speed was observed for any stability measure, indicating that the effects of shoes on running stability were similar at both running speeds. This finding does not support the hypothesis that shoe-related stability effects would become more pronounced at higher speeds. This result is consistent with the running style findings, where shoe-related biomechanical differences were likewise independent of running speed.

## Limitations

Several limitations should be considered when interpreting the findings. First, although the primary difference between the shoes was sole thickness, small differences in shoe mass were present. Although efforts were made to match shoe mass between conditions (< 50 g difference; Rodrigo-Carranza et al., 2020), potential effects of shoe mass on running biomechanics cannot be completely excluded. Second, the experiments were performed on a treadmill at constant speeds and under non-fatigued conditions. Although this controlled approach is common in running biomechanics research [9,22,63], it may not fully represent real-world running conditions. Third, the sample consisted of healthy, experienced male runners, which limits the generalizability of the results to other populations. In addition, footstrike pattern was not controlled in order to preserve natural running behavior, which may have influenced ankle joint kinematics. Fourth, gait events were identified using kinematic algorithms rather than force-based methods. Although these approaches have shown acceptable accuracy during treadmill running [51–53], small timing errors cannot be excluded. Finally, the biomechanical model represented the foot as a single rigid segment, which may reduce the accuracy of frontal-plane ankle kinematics [64].

## Conclusion

The present study investigated the effects of different shoe sole thicknesses on running style and stability during uphill running. The results indicate that AFT shoes influenced primarily vertical center-of-mass motion, vertical stiffness, and ankle joint kinematics, while spatiotemporal aspects of running style remained unchanged. Differences between the two AFT shoes were limited to frontal ankle kinematics, suggesting that sole thickness alone played a limited role in modulating running style under uphill conditions. Running stability was largely unaffected by shoes, with only a small reduction in trunk local stability observed in the moderately thick AFT shoe compared with the traditional shoe. Together, these findings indicate that both running style and running stability during uphill running were largely preserved across shoe conditions.

## Disclosure Statement

The authors report there are no competing interests to declare.

## Author Contributions

Cagla Kettner: writing – review and editing, writing – original draft, visualization, software, methodology, investigation, formal analysis, data curation, conceptualization. Bernd J. Stetter: writing – review and editing, conceptualization. Thorsten Stein: writing – review and editing, supervision, resources, project administration, methodology, funding acquisition, conceptualization.

## Acknowledgements

The authors acknowledge support from the KIT Publication Fund of the Karlsruhe Institute of Technology and the use of OpenAI’s ChatGPT (version 5.3) for language editing and readability improvement. The authors reviewed and edited all output and take full responsibility for the content.

## Funding

Adidas AG provided financial and material support for this study. The funder had no role in study design, data collection and analysis, decision to publish, or preparation of the manuscript.

## Notes

### Competing Interest Statement

The authors have declared no competing interest.

